# Long wavelength light that improves aged mitochondrial function selectively increases cytokine expression in serum and the retina

**DOI:** 10.1101/2021.11.10.468030

**Authors:** Harpreet Shinhmar, Chris Hogg, Glen Jeffery

**Affiliations:** University College London, Institute of Ophthalmology, UK

**Keywords:** inflammation, mitochondria, 670nm light, aging, photobiomodulation

## Abstract

Aged mitochondrial function can be improved with long wavelength light exposure. This reduces cellular markers of inflammation and can improve system function from fly though to human. Here, we ask what impact 670nm light has on cytokine expression using a 40 cytokine array in blood serum and retina in C57Bl6 mice. There was a relatively uniform increase in cytokine expression between 3 and 12 months of age in serum and retina.

670nm exposure was delivered daily for a week in 12 month old mice. This shifted patterns of cytokine expression in both serum and retina inducing a selective increase with some in serum increasing >5 fold. Changes in retina were smaller. In serum there were major increases in IL-7, 6, 13, 16 and 23, also in TNF-α and CXCL 9 and 10. In retina the increases were found mainly in some IL (interleukins) and CXCL’s (chemokines). A few cytokines were reduced by light exposure.

Changes in serum cytokines implies that long wavelengths impacts systemically even to unexposed tissues deep in the body. In the context of wider literature, increased cytokine expression may be protective. However, their upregulation by light merits further analysis as cytokines upregulation can also be negative.

## Introduction

Mitochondria regulate metabolism and the pace of ageing. When their membrane potential declines adenosine triphosphate (ATP) production is compromised reducing available cellular energy. This is often associated with progressive increases in reactive oxygen species (ROS) that further undermines cell function^1^. However, long wavelength light (650-900nm) is able to reverse many of these features in ageing and disease, particularly in the CNS with its high metabolic demand and mitochondrial dependence^2^.

Exposure to longer wavelengths has been widely shown to have therapeutic value in the CNS. In invertebrates it extends lifespan and improves aged motor skills, cognition and visual function^3–6^. In mammals it has similar impact and also reduces cellular markers of inflammation and the pace of age related cell loss^7–9^. Its application is now extended to humans where aged visual function is significantly improved^10^.

There is limited evidence that longer wavelength exposure also impacts on immunity^11–13^. However, this remains relatively unexplored. A key question here relates to whether changes in immunity induced by longer wavelengths can also be found in serum. If this were the case, then it may imply that the impact of such lights can act systemically. There has been evidence for this when longer wavelengths have been targeted at distal regions of the body and have had positive impacts on the retina^13^. But the mechanism for this has not been revealed.

The absence of data on the interactions between immunity and longer wavelengths is problematic for the development of therapies based on their use. In this study we run cytokine arrays on mice revealing age related changes in the retina and in serum. We then expose aged mice to a commonly used long wavelength, 670nm, and assess its impact on the cytokine expression. The hypothesis is that consistent with improved mitochondrial function, there will be a decline in the patterns of cytokine expression following 670nm light exposure. This proved not to be the case.

## Methods

### Animals

C57 mice were used throughout with full local and UK government approval. Mice were used at two ages, young at 3 months (N= 6) and old at 12 months (N=6). Two experiments were undertaken. First, young (N= 6) and old mice (N=6) were compared for cytokine expression in serum and retina. Second, old mice (N=6, 12 months old) were exposed to 670nm (40mW/cm2) daily at 10am for 15 mins for 7 days and compared against aged controls (N=6). All mice were killed by cervical dislocation. Eyes were rapidly removed, and the retina extracted on ice and processed as tissue lysates as below. Bloods were taken via cardiac puncture. Retinal and serum samples were pooled within groups.

### Blood Serum Collection

Blood was rapidly harvested into tubes and allowed to coagulate on ice for approximately 20 minutes. After the blood had coagulated for 20 minutes the tubes were then centrifuged at 2000 × g for 15 minutes and the serum was transferred to a new tube and snap frozen at −80°c; this serum was used for cytokine arrays. Protein concentration was calculated using a BCA Assay kit (Thermo Scientific). Serum was pooled per group so that 150μl could be added to each cytokine array membrane.

### Tissue Lysates

The retinal samples were homogenized with a protease cocktail inhibitor mix. Samples were then frozen with the addition of Triton X-100 to a final concentration of 1%. Lysates were thawed and centrifuged at 10,000 × g for 10 minutes and the resulting supernatant collected. Protein concentration was calculated using a BCA Assay (Thermo Scientific). Samples were pooled per group so that a final concentration of 200μg of protein could be added to each membrane.

### Cytokines

Cytokine levels were assessed using the Proteome Profiler Mouse Cytokine Array Panel A (R & D Systems, Minneapolis, MN, USA) according to the protocol below. Array nitrocellulose membranes were incubated for 1 hour on a rocking platform with Array buffer 6, a buffered protein base with preservatives, which served as a blocking buffer. Whilst blocking, the samples (blood serum and retina) were incubated for 1 hour with Array buffer 4, a buffered protein base with preservatives, and a biotinylated antibody cocktail. The membranes were then incubated overnight on the rocking platform with the sample/antibody mixture. The following day the membranes were washed several times with a wash buffer, a solution of buffered surfactant. The arrays were then incubated for 30 minutes with Streptavidin-HRP (Streptavidin conjugated to horseradish-peroxidase) on a rocking platform. The membranes were then washed again several times with wash buffer. Any excess liquid was drained off the membranes before a Chemi reagent mix, made up of stabilised hydrogen peroxidase and luminol, was applied and incubated for 1 minute. Any excess reagent was removed before the membranes were fixed in an autoradiography film cassette and exposed to an X-ray film. Protein Array Analyzer for Image J was used to quantify and determine spot density from the X-ray film.

## Results

Cytokine expression was compared across young and old mice in serum and in the retina. In both cases the data are displayed in the figures with the highest concentration to the left running to the lowest on the right.

Profiles in serum showed that for each of the 40 cytokines measure levels were greater in older mice than in the younger cohort (Figure 1). Differences between the two groups over the 40 cytokines were approximately 36.5%, but differences were not even. Hence, small changes were found in IL-2, 3, 5, 6,10 large differences were present in CXCL12, CCL12. The heat map below the histograms gives the proportionate changes in each cytokine.

**Figure 1:**
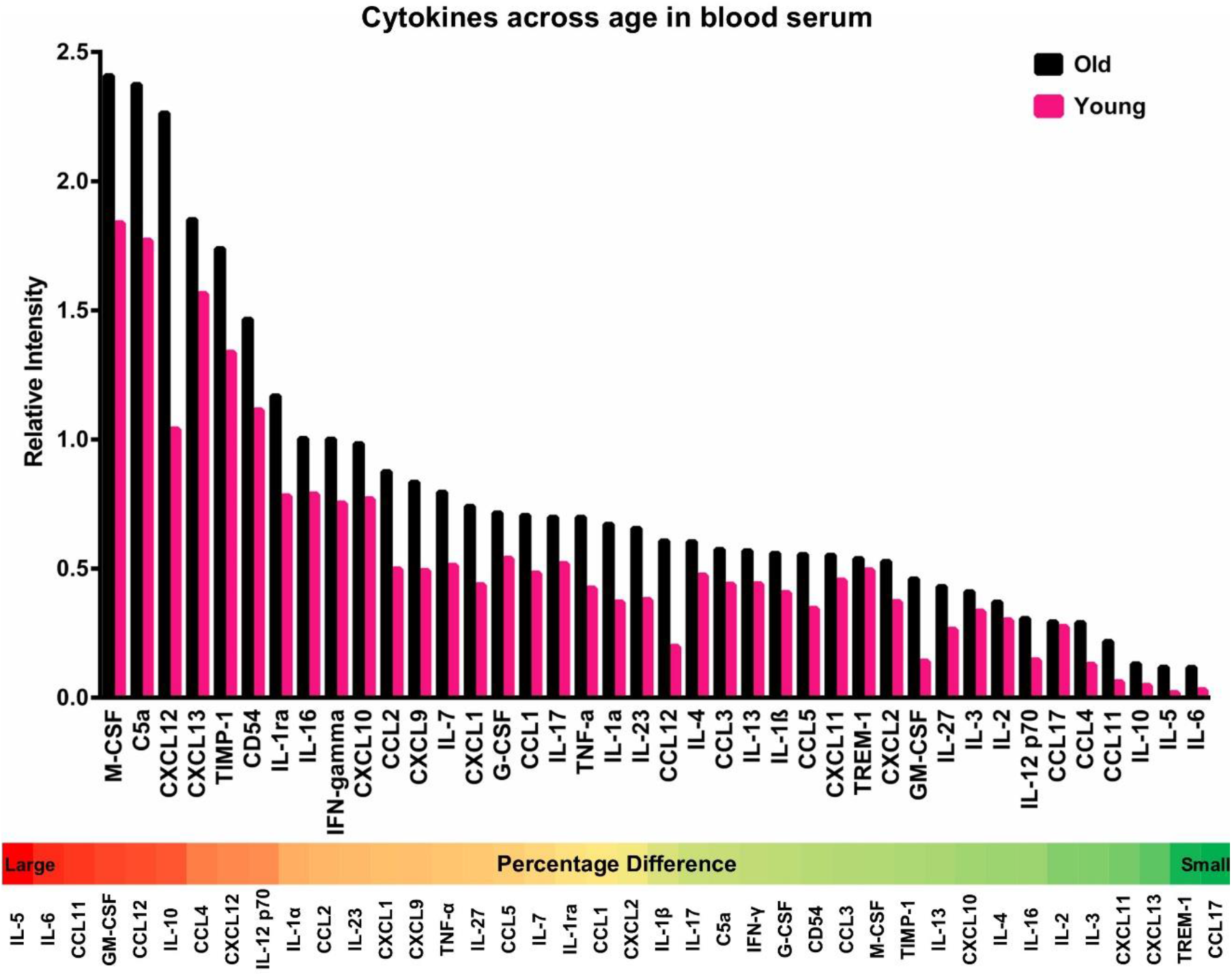
The pattern of cytokine expression with age in the blood serum of young 3 month old (N=6. Pink) and 12 month old (N=6. Black) C57Bll6 mice. Data were pooled across mice at both ages. All 40 cytokines displayed an increase with age in serum. The overall aged increase was 36.5%. The upper graph has the cytokines ordered with the highest expressing on the left and the lowest expressing on the right. Below is a heat map showing the relative percentage increase in expression of each of the cytokines with red indicating a large increase and green a small increase.

Cytokine expression patterns across young and old mice in the retina were largely similar to that found in serum, with all but one cytokine expressed at higher levels in old mice compared to young (Figure 2). However, differences between the age groups were not as marked in retina compared to serum, with an approximate difference of 26%. The only exception was CXCL13, which is active on mature lymphocytes and is associated with the generation of broadly neutralising antibodies^14,15^. Again, the data are shown as a heat map below showing relative change between the two age groups.

**Figure 2:**
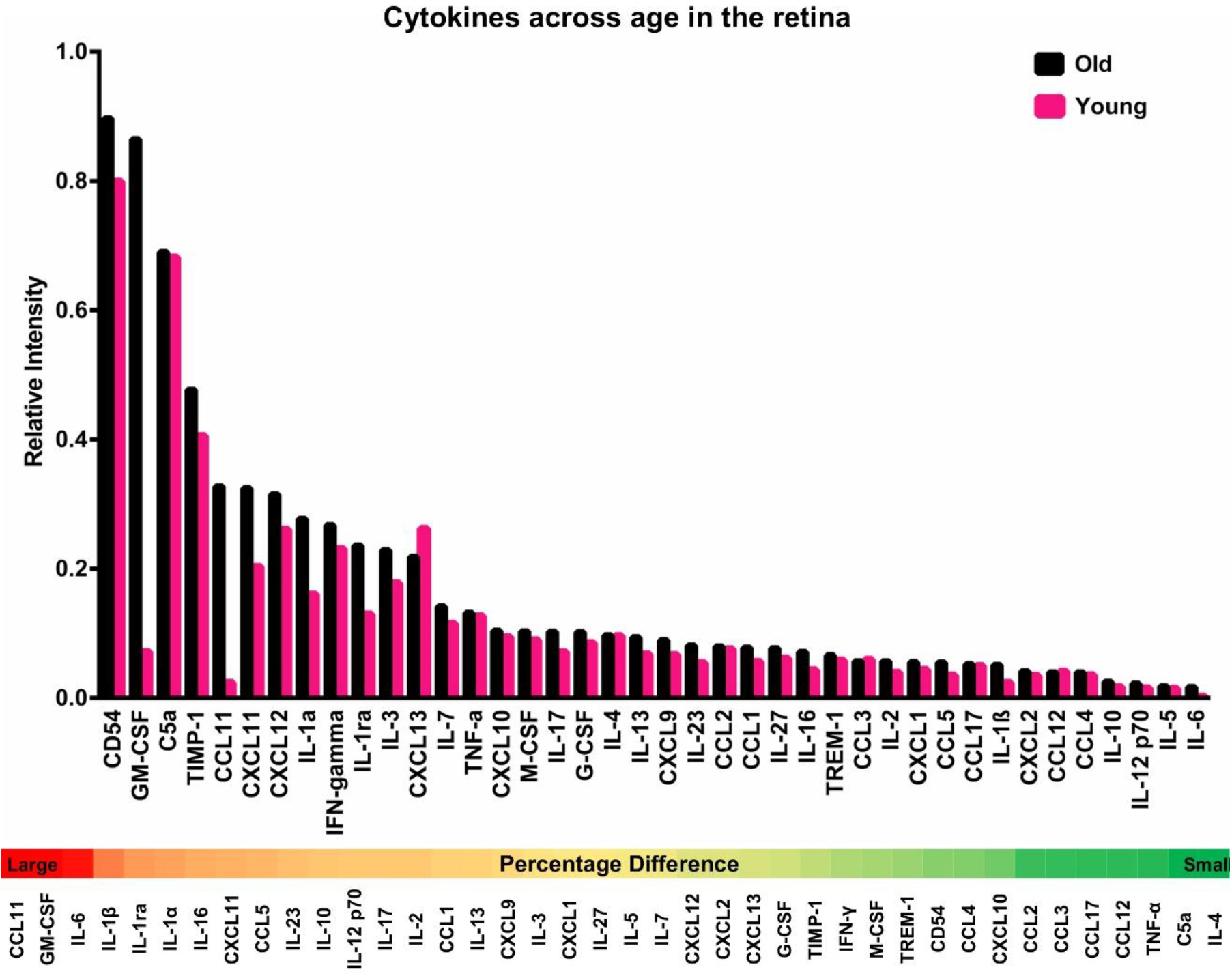
The pattern of cytokines with age in the retina of young 3 month old (N=6. Pink) and 12 month old (N= 6. Black) C57Bll6 mice. Data were pooled within groups. The mice used are the same as those in Figure 1 used for serum. 39 out of 40 cytokines displayed an increase with age in the retina, with the exception of CXCL13. The average increase in cytokine expression between young and old animals was 26%. The upper graph has the cytokines ordered with the highest expressing on the left and the lowest expressing on the right. Below is a heat map showing the relative percentage increase in expression of each of the cytokines with red indicating a large increase and green a small increase.

12 month old animals were examined after exposure to 670nm light and compare against unexposed controls. Here the results were different from that found across the two untreated age groups. In both serum (Figure 3) and retina (Figure 4) some cytokines were reduced in expression following 670nm exposure, although reductions in expression were less frequent than increases.

**Figure 3:**
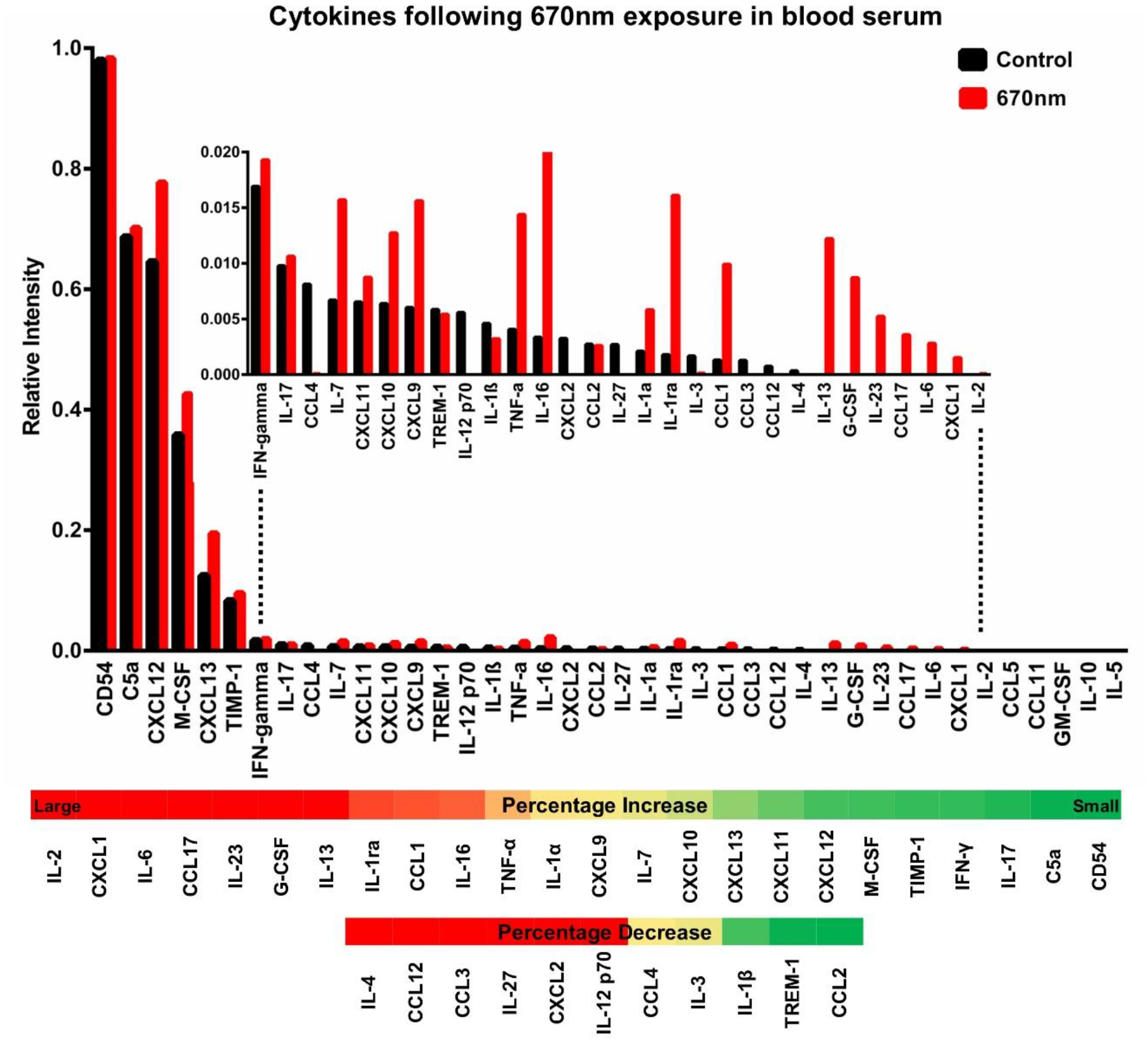
Pattern of cytokines in the blood serum of 12-month old C57Bll6 mice following exposure with 670nm (N=6. Red) compared to aged match controls (N=6. Black). Data were pooled within groups. The Y-axis has been expanded in the inset to further monitor the differences that were on a smaller scale. In both cases cytokines are ordered with the highest expressing on the left and the lowest expressing on the right. Below is a heat map showing the relative percentage increase or decrease in expression of each of the cytokines with red indicating a relatively large change and green a relatively small change.

**Figure 4:**
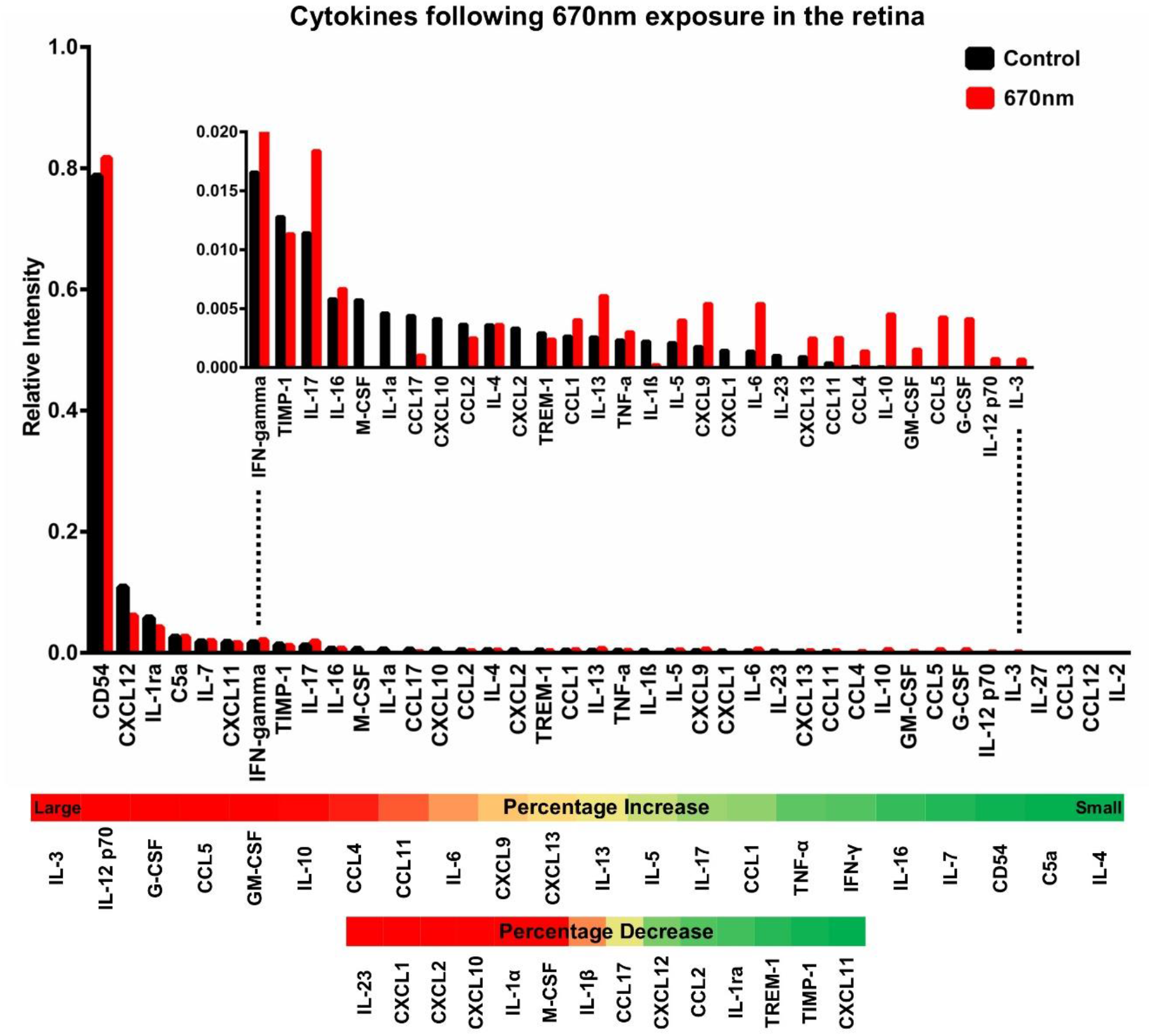
Pattern of cytokines in the retina of 12-month old C57Bl6 mice following exposure with 670nm (N=6. Red) compared to aged match controls (N=6. Black). The mice used are the same as those for serum in figure 3. Data were pooled within groups. The Y-axis has been expanded in the inset to further monitor the differences that were on a smaller scale. In both cases cytokines are ordered with the highest expressing on the left and the lowest expressing on the right. Below is a heat map showing the relative percentage increase or decrease in expression of each of the cytokines with red indicating a relatively large change and green a relatively small change.

In serum the proportional increases appeared to be large, there were marked increases in IL-2, 6, 13 and 23, along with CXCL1 and marked decreases in IL-4 CCL3 and 12. Hence, within interleukins IL’s there were both increases and decreases.

In retina changes following 670nm exposure were less marked than in serum. Further the patterns of change in retina were different from those found in serum. Hence there were marked increases in IL-3 and 10, along with G-CSF and marked decreases in IL-3, CXCL1, 2 and 10.

## Discussion

Cytokine expression in serum and in the retina increases with age and with exposure to 670nm light. Aged increases are relatively uniform and no cytokine was found to decrease in the older age group compared to the young. The increase in cytokine expression was unexpected following 670nm exposure. Increases particularly in serum were marked and specific. While some increased by as much as 5 fold others did not change and some were reduced.

The parameters we have used in terms of the mice, their age and patterns of 670nm exposure, in duration and energy, are consistent with our previous studies. These have consistently shown improvements in mitochondrial function along with reduced cell death and improved retinal function^9,16–18^. In the retina they have also shown reduced markers of tissue inflammation and stress using immune staining on tissue sections^7,19,20^. But commonly they have not been the same inflammatory markers that have been used here, rather they have often been those associated with complement^7,8^.

In retinae reductions have been shown in TNF-α and complement proteins in immune staining^19,20^. The key difference between these results and those presented in is study is that here the cytokines, particularly those in the serum, were unbound. It is possible that an overall less marked increase in cytokine expression in the retina after 670nm exposure compared to serum is because this includes bound and unbound elements, and perhaps the bound elements were reduced. Hence TNF-α as expressed in tissue section and immune staining is significantly reduced following 670nm exposure^7,8^, but is elevated in serum in this study using identical exposure protocols. It did not appear to change much in the array on the retina after 670nm exposure but this picture may be blurred by inclusion of bound and unbound elements in retinal samples.

If there is consistency between previous studies and the data presented here, it highlights that the selective cytokines that are up regulated are likely to be protective. There is evidence for the protective impact of cytokines^21,22^ and often different cytokines act in complex synergy^23^. The consequence is that it is very difficult to determine what the impact of up and down regulation of any cytokine is as it does not occur in isolation but is likely to be highly interactive with the dynamics of other cytokines and inflammatory agents. Hence, while those that shift in either direction following 670nm exposure are associated with improved cellular and organ function determining which is doing what is beyond the scope of this study.

Our data do offer a potential explanation for enigmatic results that have been described as the abscopal effect when treatment of tissues distal to pathology is found to be effective. Two labs have shown that retinal pathology may be partially ameliorated by long wavelength exposure to regions other than the eye^13,24^, while others have shown similar general systemic effects following focal long wavelength exposure^25,26^. Our results provide a potential route for this effect via shifts in the patterns of circulating cytokines.

This study has used experimental metrics consistent with those employed previously for consistency. However, there is far less data available for potential interactions between light, mitochondria and cytokines. The relationship between immunity and mitochondrial function is far from clea^27–29^. Consequently, we have little idea of whether this snapshot we provide at one time following a week’s exposure to 670nm light is representative. It is possible that cytokine expression is reduced after 670nm light exposure and that what we are viewing is actually a rebound effect. ATP upregulation following a single 1 min 670nm exposure occurs within an hour and improves respiration significantly for almost a week^3,11^. But we know no more about the dynamics of this let alone how immunity changes over time. To have used different time points/exposures would have robbed our data set from its relationship to the majority of other 670nm studies undertaken on mice that reveals positive effects. However, to explore the large number of permutations and combinations of lights and potential post exposure time periods would be a larger undertaking than possible here.

The use of mice represents an additional problem. The impact of long wavelength light appears to be species independent, even having commonality between invertebrates and mammals including humans^4,10,18^. However, mouse immunity is fundamentally different from that of humans, which is exacerbated by laboratory mice being maintained in pathogen free environments^30–33^. When cytokine arrays specific for mice and human are used in retinae from mice and old world primates in ageing, the patterns of up regulation in the two species have almost no relationship with one another^34^. Hence while our results have a degree of significance, the problematic nature of mouse immunity needs to be kept in mind.

While long wavelength light has shown largely consistent patterns of improvement from cells through to system function in ageing, the fact that it is associated with increased cytokine expression remains an issue. It is possible that this is protective and that cytokine expression is different at separate time points following exposure. But it is clearly important that the experiments undertaken here, particularly those on serum, are repeated in aged humans where long wavelengths are increasingly being employed in ageing and age related disease.

## Acknowledgments

We thank Jaimie Hoh Kam and Mike Powner for their comments on the manuscript

## Author contributions

HS and GJ designed the experiment. HS performed the experiment. GJ and HS wrote the manuscript. CH provided equipment. All authors reviewed the manuscript.

## Funding

BBSRC BB/N000250/1

## Competing interests

The authors declare no competing interests

## Notes

### Competing Interest Statement

The authors have declared no competing interest.

## References

1. López-Otín, C., Blasco, M. A., Partridge, L., Serrano, M. & Kroemer, G. The hallmarks of aging. Cell 153, 1194 (2013).

2. Mitrofanis, J. & Jeffery, G. Does photobiomodulation influence ageing? Aging (Albany. NY). 10, 2224–2225 (2018).

3. Weinrich, T. W., Hogg, C. & Jeffery, G. The temporal sequence of improved mitochondrial function on the dynamics of respiration, mobility, and cognition in aged Drosophila. Neurobiol. Aging 70, 140–147 (2018).

4. Weinrich, T. W., Coyne, A., Salt, T. E., Hogg, C. & Jeffery, G. Improving mitochondrial function significantly reduces metabolic, visual, motor and cognitive decline in aged Drosophila melanogaster. Neurobiol. Aging 60, 34–43 (2017).

5. Michael, B. P., Thomas, E. S., Chris, H. & Glen, J. Improving mitochondrial function protects bumblebees from neonicotinoid pesticides. PLoS One 11, 1–11 (2016).

6. Begum, R. et al. Near-infrared light increases ATP, extends lifespan and improves mobility in aged Drosophila melanogaster. Biol. Lett. 11, 20150073–20150073 (2015).

7. Begum, R., Powner, M. B., Hudson, N., Hogg, C. & Jeffery, G. Treatment with 670 nm Light Up Regulates Cytochrome C Oxidase Expression and Reduces Inflammation in an Age-Related Macular Degeneration Model. PLoS One 8, 1–11 (2013).

8. Kokkinopoulos, I., Colman, A., Hogg, C., Heckenlively, J. & Jeffery, G. Age-related retinal inflammation is reduced by 670 nm light via increased mitochondrial membrane potential. Neurobiol. Aging 34, 602–609 (2013).

9. Sivapathasuntharam, C., Sivaprasad, S., Hogg, C. & Jeffery, G. Improving mitochondrial function significantly reduces the rate of age related photoreceptor loss. Exp. Eye Res. 185, 107691 (2019).

10. Shinhmar, H. et al. Optically Improved Mitochondrial Function Redeems Aged Human Visual Decline. Journals Gerontol. – Ser. A Biol. Sci. Med. Sci. 75, e49–e52 (2020).

11. Powner, M. B., Priestley, G., Hogg, C. & Jeffery, G. Improved mitochondrial function corrects immunodeficiency and impaired respiration in neonicotinoid exposed bumblebees. PLoS One 16, 36–38 (2021).

12. Golovynska, I. et al. Macrophages Modulated by Red/NIR Light: Phagocytosis, Cytokines, Mitochondrial Activity, Ca 2+ Influx, Membrane Depolarization and Viability. Photochem. Photobiol. (2021). doi:10.1111/php.13526

13. Cheng, Y. et al. Photobiomodulation inhibits long-term structural and functional lesions of diabetic retinopathy. Diabetes 67, 291–298 (2018).

14. Kazanietz, M. G., Durando, M. & Cooke, M. CXCL13 and its receptor CXCR5 in cancer: Inflammation, immune response, and beyond. Front. Endocrinol. (Lausanne). 10, 1–15 (2019).

15. Havenar-Daughton, C. et al. CXCL13 is a plasma biomarker of germinal center activity. Proc. Natl. Acad. Sci. U. S. A. 113, 2702–2707 (2016).

16. Sivapathasuntharam, C., Sivaprasad, S., Hogg, C. & Jeffery, G. Aging retinal function is improved by near infrared light (670 nm) that is associated with corrected mitochondrial decline. Neurobiol. Aging 52, 66–70 (2017).

17. Kaynezhad, P., Tachtsidis, I. & Jeffery, G. Optical monitoring of retinal respiration in real time: 670 nm light increases the redox state of mitochondria. Exp. Eye Res. 152, 88–93 (2016).

18. Gkotsi, D. et al. Recharging mitochondrial batteries in old eyes. Near infra-red increases ATP. Exp. Eye Res. 122, 50–53 (2014).

19. Hoh Kam, J., Lenassi, E., Malik, T. H., Pickering, M. C. & Jeffery, G. Complement component C3 plays a critical role in protecting the aging retina in a murine model of age-related macular degeneration. Am. J. Pathol. 183, 480–492 (2013).

20. Hoh Kam, J., Lynch, A., Begum, R., Cunea, A. & Jeffery, G. Topical cyclodextrin reduces amyloid beta and inflammation improving retinal function in ageing mice. Exp. Eye Res. 135, 59–66 (2015).

21. Lilic, D. et al. Deregulated production of protective cytokines in response to Candida albicans infection in patients with chronic mucocutaneous candidiasis. Infect. Immun. 71, 5690–5699 (2003).

22. Cai, J. et al. Protective/reparative cytokines are suppressed at high injury severity in human trauma. Trauma Surg. Acute Care Open 6, e000619 (2021).

23. Chabaud, M., Page, G. & Miossec, P. Enhancing Effect of IL-1, IL-17, and TNF-α on Macrophage Inflammatory Protein-3α Production in Rheumatoid Arthritis: Regulation by Soluble Receptors and Th2 Cytokines. J. Immunol. 167, 6015–6020 (2001).

24. Johnstone, D. M. et al. Indirect application of near infrared light induces neuroprotection in a mouse model of parkinsonism – An abscopal neuroprotective effect. Neuroscience 274, 93–101 (2014).

25. Rochkind, S. et al. Systemic effects of low‐power laser irradiation on the peripheral and central nervous system, cutaneous wounds, and burns. Lasers Surg. Med. 9, 174–182 (1989).

26. Rodrigo, S. M. et al. Analysis of the systemic effect of red and infrared laser therapy on wound repair. Photomed. Laser Surg. 27, 929–935 (2009).

27. Missiroli, S. et al. The role of mitochondria in inflammation: From cancer to neurodegenerative disorders. J. Clin. Med. 9, (2020).

28. Hoffmann, M. B. et al. Minor effect of blue-light filtering on multifocal electroretinograms. J. Cataract Refract. Surg. 36, 1692–1699 (2010).

29. Qualls, A. E., Southern, W. M. & Call, J. A. Mitochondria-cytokine crosstalk following skeletal muscle injury and disuse: A mini-review. Am. J. Physiol. – Cell Physiol. 320, C681–C688 (2021).

30. Mestas, J. & Hughes, C. C. W. Of Mice and Not Men: Differences between Mouse and Human Immunology. J. Immunol. 172, 2731–2738 (2004).

31. Willyard, C. Squeaky clean mice could be ruining research. Nature 556, 16–18 (2018).

32. Abolins, S. et al. The comparative immunology of wild and laboratory mice, Mus musculus domesticus. Nat. Commun. 8, 1–13 (2017).

33. Sellers, R. S. Translating Mouse Models: Immune Variation and Efficacy Testing. Toxicol. Pathol. 45, 134–145 (2017).

34. Kam, J. H. et al. Fundamental differences in patterns of retinal ageing between primates and mice. Sci. Rep. 9, 1–14 (2019).

